# Human-specific staphylococcal virulence factors enhance pathogenicity in a humanised zebrafish C5a receptor model

**DOI:** 10.1101/2020.02.18.955021

**Authors:** Kyle D. Buchan, Michiel van Gent, Tomasz K. Prajsnar, Nikolay V. Ogryzko, Nienke W.M. de Jong, Julia Kolata, Simon J. Foster, Jos A.G. van Strijp, Stephen A. Renshaw

## Abstract

*Staphylococcus aureus* infects approximately 30% of the human population and causes a spectrum of pathologies ranging from mild skin infections to life-threatening invasive diseases. The strict host specificity of its virulence factors has severely limited the accuracy of *in vivo* models for the development of vaccines and therapeutics. To resolve this, we generated a humanised zebrafish model and determined that neutrophil-specific expression of the human C5a receptor conferred susceptibility to the *S. aureus* toxins PVL and HlgCB, leading to reduced neutrophil numbers at the site of infection and increased infection-associated mortality as a direct result of the interaction between *S. aureus* and the receptor. These results show that humanised zebrafish provide a valuable platform to study the contribution of human-specific *S. aureus* virulence factors to infection *in vivo* that could facilitate the development of novel therapeutic approaches and essential vaccines.

## Introduction

*Staphylococcus aureus* is a highly specialised pathogen that colonises 30% of the human population and causes a variety of mild to severe illnesses ranging from skin and soft-tissue infections to necrotising pneumonia, endocarditis and septicaemia (*1*). In the USA, as many as 50% of *S. aureus* infections are caused by antibiotic-resistant strains (*2*), with methicillin-resistant *S. aureus* (MRSA) among the leading causes of death by a single infectious agent (*3*), emphasising the need for development of alternative therapies or vaccines. Despite promising results from vaccine studies utilising bacterial surface components and toxins as antigens, these candidates have largely failed to translate from traditional infection models to humans (*4*). A likely reason for this is the inability of current *in vivo* models to accurately recapitulate human infections, as *S. aureus* expresses a variety of strictly human-specific virulence factors that are ineffective in these models (*4*). Although their contribution to natural infection remains poorly understood, these virulence factors broadly target the host innate immune system to impair the complement system, oxidative enzymes, chemotactic proteins and phagocytic cells in order to evade recognition and destruction (*5*). To properly recapitulate human infections and to study the contributions of *S. aureus* virulence factors to infectivity and pathogenesis *in vivo*, a new humanised infection model is required.

Three *S. aureus* virulence factors interact with the human C5a receptor (C5AR1, CD88), a seven-transmembrane loop G-protein coupled receptor (GPCR) that is highly expressed on the surface of neutrophils (*6*) and recognises host anaphylatoxin C5a released during complement activation to control phagocyte activation and chemotaxis. The targeting of C5AR1 by multiple *S. aureus* virulence factors appears to be specifically focused against neutrophils, which form an essential line of defence against staphylococcal infection (*7*). The two bicomponent pore-forming toxins Panton-Valentine Leukocidin (PVL) and γ-Haemolysin CB (HlgCB) target C5AR1 to recognise and lyse phagocytic cells by forming β-barrel pores in the cell membrane (*8, 9*). The toxins are secreted as two subunits, known as the S- (slow, LukS-PV/HlgC) and F- (fast, LukF-PV/HlgB) subunits due to their chromatography elution profiles. Besides inducing cell lysis, S-subunits also disrupt chemotaxis by competitively inhibiting receptor signalling (*8, 9*). In addition, the chemotaxis inhibitory protein of *S. aureus* (CHIPS) is a 14 kDa protein that prevents C5a-mediated chemotaxis by binding directly to the N-terminus of C5AR1 (*10*). The high-affinity protein-protein interactions between virulence factors and C5AR1, which have been characterised at the amino-acid level, are highly human specific and consequently our insight into the roles they play during infection is limited due to a lack of suitable humanised infection models (*4, 11*).

The zebrafish (*Danio rerio*) is a widely used model organism for investigating bacterial infections and the innate immune system (*12, 13*). Due to their optical transparency, ability to produce high numbers of offspring and genetic tractability, zebrafish offer many unique approaches over existing infection models. Zebrafish have a fully-developed innate immune system by two days post-fertilisation characterised by the presence of mature phagocytic cells (*14*) and a complement system that is highly homologous to humans (*15*).

In this study, we developed a humanised C5AR1 knock-in zebrafish infection model and determined the contribution of the *S. aureus* toxins PVL and HlgCB to infection *in vivo*. Whereas wild type zebrafish neutrophils were resistant to toxin-mediated lysis, we show that neutrophil-specific C5AR1 expression confers sensitivity of zebrafish neutrophils to PVL and HlgCB-mediated lysis *in vivo*. Humanised zebrafish displayed reduced neutrophil abundance at the sites of infection and increased *S. aureus*-associated mortality as a result of the direct interaction between *S. aureus* and the human C5a receptor when expressed by zebrafish neutrophils. In conclusion, our studies not only illustrate the critical contribution of PVL and HlgCB to *in vivo* infection and pathogenesis, but also show the significance of humanised zebrafish as a novel platform to investigate the activities of human-specific virulence factors *in vivo* and to accurately recapitulate natural human infection in a model organism.

## Results

### Zebrafish possess a functional C5aR that is responsive to serum-derived C5a

To study the C5a-C5aR signalling axis in zebrafish, we first expressed the zebrafish C5a receptor (*c5ar1*) in the human monocyte-like cell line U937 (U937-dreC5aR) by lentiviral transduction and measured the receptor’s ability to bind and respond to recombinant zebrafish and human C5a (dreC5a and hsaC5a, respectively) using previously established methods (*8*). As controls, we generated cells stably expressing the human C5a receptor (U937-hsaC5aR), or an empty vector control (U937-EV). Firstly, flow-cytometric analysis of recombinant, FITC-labelled C5a-binding capacities showed that U937-dreC5aR cells specifically bound dreC5a, but not hsaC5a (**Fig. 1A)**. Conversely, U937-hsaC5aR cells strongly bound hsaC5a and only minimally interacted with dreC5a, suggesting that both receptors bind C5a in a species-specific manner. Activation of GPCRs, including the C5a receptor, results in the induction of intracellular signalling cascades culminating in the cytosolic release of intracellular calcium stores (*16*). Accordingly, we evaluated the signalling ability of the receptors by measuring intracellular Ca^2+^ release following receptor stimulation using the Fluo-3AM probe. Treatment of U937-dreC5aR cells with recombinant dreC5a provoked robust calcium release, indicating successful receptor ligation and activation, whereas U937-hsaC5aR expressing cells responded minimally to dreC5a (**Fig. 1B**). Importantly, zymosan-activated zebrafish serum induced a response similar to recombinant C5a, indicating that the zebrafish C5aR responds to physiological concentrations of zebrafish C5a in activated fish serum under these conditions (**Fig. 1B**). Conversely, U937-hsaC5aR cells responded robustly to hsaC5a or human activated serum treatment, but not dreC5a treatment. Taken together, these data show that the zebrafish *c5ar1* gene encodes a functional surface receptor that interacts with, and is activated by, physiological concentrations of C5a. Furthermore, the human and zebrafish C5a receptor and ligand pairs are species specific and are not interchangeable.

**Figure 1.**
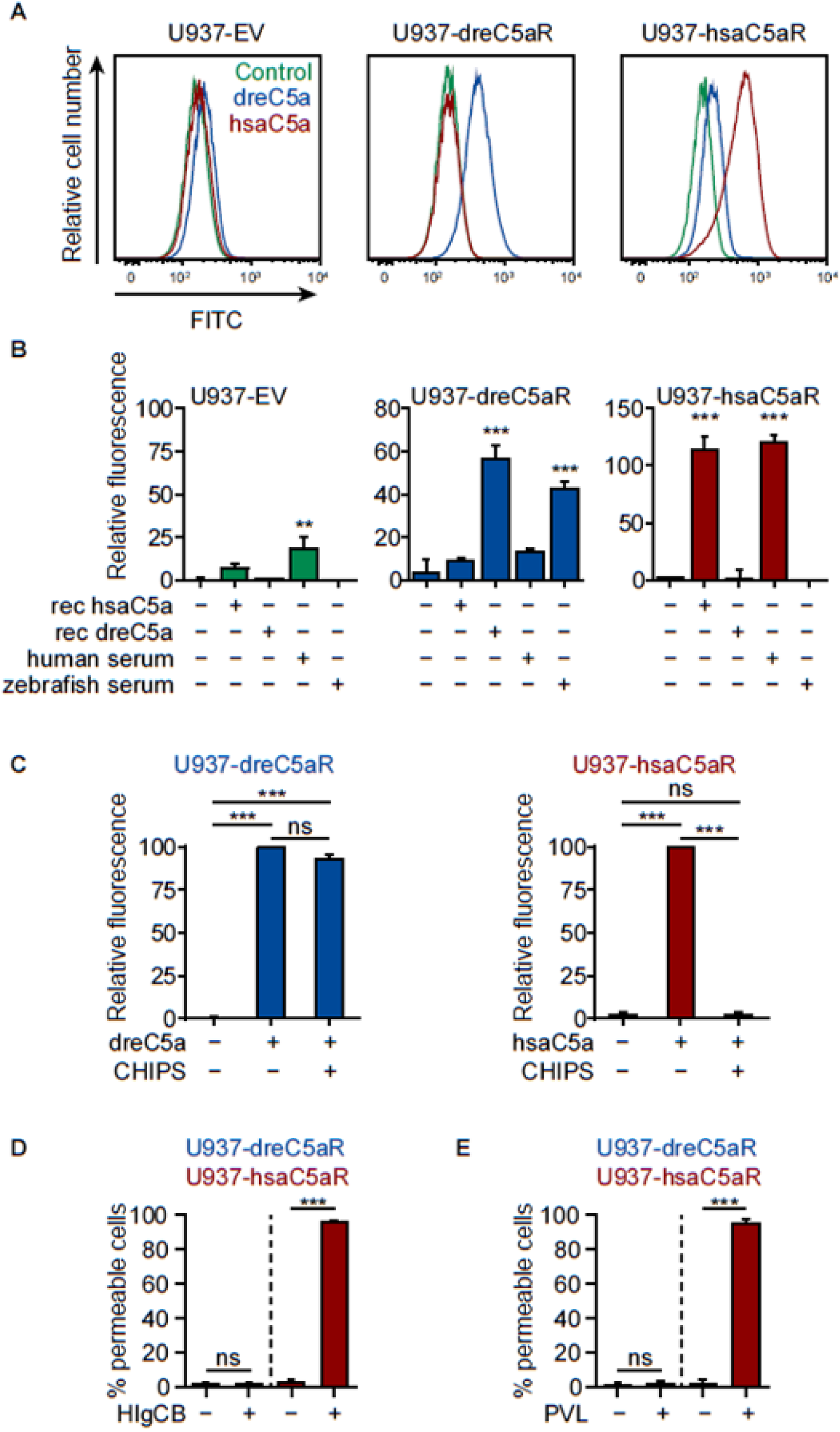
Zebrafish possess a functional C5aR that is insensitive to targeting by *S. aureus* virulence factors CHIPS, PVL, and HlgCB. **A)** Binding of FITC-labelled zebrafish C5a (dreC5a) or human C5a (hsaC5a) to U937 cells stably expressing the zebrafish C5a receptor (dreC5aR), the human C5a receptor (hsaC5aR) or an empty vector control (EV) was determined by flow cytometry. **B)** Relative C5aR activity following treatment with recombinant human or zebrafish C5a or zymosan-activated serum was determined by measuring cytosolic calcium release using a Ca^2+^-sensitive fluorescent probe (Fluo-3AM) and displayed as a percentage relative to the positive control (10 µM ionomycin treatment, set at 100%). **C)** C5aR activity following treatment with recombinant human or zebrafish C5a with or without 10 μg/ml recombinant CHIPS was determined by measuring cytosolic Ca^2+^ release using a Ca^2+^-sensitive fluorescent probe (Fluo-3AM) by flow cytometry and displayed as mean percentage ± SD relative to the C5a-treated sample without CHIPS (middle bar, set at 100%). **D,E)** Pore formation in U937-dreC5aR (blue) or U937-hsaC5aR (red) cells following 30 min incubation with 10 μg/ml HlgCB (D) or 10 μg/ml PVL (E), as measured by percentage DAPI-positive cells by flow cytometry. Data are presented as means ± SD; ns, not significant; *, *p*<0.05; **, *p*<0.01; ***, *p*<0.001; two-way ANOVA with Dunnett’s multiple comparisons correction of each sample versus the untreated control (A-C) or *t*-test (D,E).

### Zebrafish C5aR is resistant to human-specific virulence factors

The three *S. aureus* C5AR1-targeting virulence factors, CHIPS, PVL, and HlgCB, are known to display strict human specificity and are unable to interact with C5aRs expressed by several other species (*8, 9*). However, their ability to target zebrafish complement components is unknown. To test this, we first assessed the functionality of the zebrafish C5a receptor in the absence and presence of the inhibitory protein CHIPS. We observed that U937-dreC5aR cells retained complete responsiveness to dreC5a in the presence of CHIPS at concentrations that completely inhibited U937-hsaC5aR cells, indicating that CHIPS is ineffective at targeting the zebrafish receptor (**Fig. 1C**). Based on our detailed molecular understanding of the interactions between CHIPS and the 21-amino acid binding site in the N-terminus of the human C5aR, we predicted that a few amino-acid changes may be sufficient to render the zebrafish C5aR sensitive to the inhibitory actions of CHIPS. To test this, we generated ten rationally designed variants of the zebrafish receptor, substituting human residues at key points within the CHIPS-binding site (*17*). All ten variants showed normal surface expression and endogenous signalling activity in response to recombinant dreC5a similar to the wildtype zebrafish receptor (**Fig. S1**). Notably, activation of three partly humanised variants (**Fig. S2, variants A, H and I**) by dreC5a was effectively inhibited in the presence of CHIPS, the most conservative change requiring only three amino acid substitutions (**Fig. S2, variant H**). These data indicated that sensitisation of dreC5aR to CHIPS can be achieved with only three amino acid changes in the endogenous receptor.

Next, we determined the sensitivity of zebrafish C5aR-expressing cells to pore-formation and lysis by the *S. aureus* toxins PVL and HlgCB. Whereas U937-hsaC5aR cells were efficiently permeabilised in the presence of the recombinant leukocidin components of PVL or HlgCB, U937-dreC5aR cells were resistant to these same concentrations of PVL and HlgCB (**Fig. 1D,E**). We then aimed to identify the minimal amino-acid changes in the zebrafish C5aR that are sufficient to gain sensitivity to PVL and/or HlgCB using the same strategy as we used to gain CHIPS sensitivity. We tested an extensive set of dreC5aR variants with many combinations of (partly) humanised intracellular and extracellular loops and individual amino acids that are known to be involved in the PVL/HlgCB interaction with the hsaC5aR. Unfortunately, we failed to achieve toxin-sensitivity while maintaining surface expression and signalling capability of the dreC5aR. In conclusion, we found that the wild type zebrafish C5aR is insensitive to the lytic activity of *S. aureus* virulence factors CHIPS, PVL, and HlgCB at the concentrations tested. While CHIPS sensitivity can be achieved with only three amino-acid changes in dreC5aR, we were unable to identify dreC5aR variants that gained PVL and/or HlgCB sensitivity.

### Generation of transgenic zebrafish with neutrophil specific human C5AR1 expression

Next, we sought to develop an *in vivo* infection model to study the role of *S. aureus* pore-forming toxins during natural infection. Because we were unable to generate a humanised drC5aR sensitive to PVL and/or HlgCB *in vitro*, we instead introduced the complete human C5a receptor into zebrafish neutrophils. To this end, a transgenic construct directing expression of a fluorescent, Clover-tagged C5AR1 driven by the zebrafish neutrophil-specific *lyz* promoter (*18*) was introduced into the zebrafish genome by Tol2 transgenesis (*19*), producing the transgenic line *Tg(lyz:hsaC5AR1-Clover)sh505*. To verify whether C5AR1-Clover expression was restricted to zebrafish neutrophils, we crossed this line to the established transgenic zebrafish line *Tg(lyz:nfsB-mCherry)sh260* that displays neutrophil-specific mCherry expression (*18*). In the double-transgenic larvae, we observed Clover expression exclusively in the mCherry-positive cells, confirming that the C5AR1 protein is expressed specifically in zebrafish neutrophils (**Fig. 2A,B**). Furthermore, whereas mCherry showed general cytoplasmic localisation, the C5AR1-associated Clover signal was enriched at the cell membrane of neutrophils from *Tg(lyz:hsaC5aR1-Clover)sh505* zebrafish, suggesting that the cell-surface expression of C5AR1 observed in human neutrophils is correctly recapitulated in the humanised zebrafish system (**Fig. 2C,D**). Notably, the total number of neutrophils in these fish was unaffected by transgene expression (**Fig. S3**).

**Figure 2.**
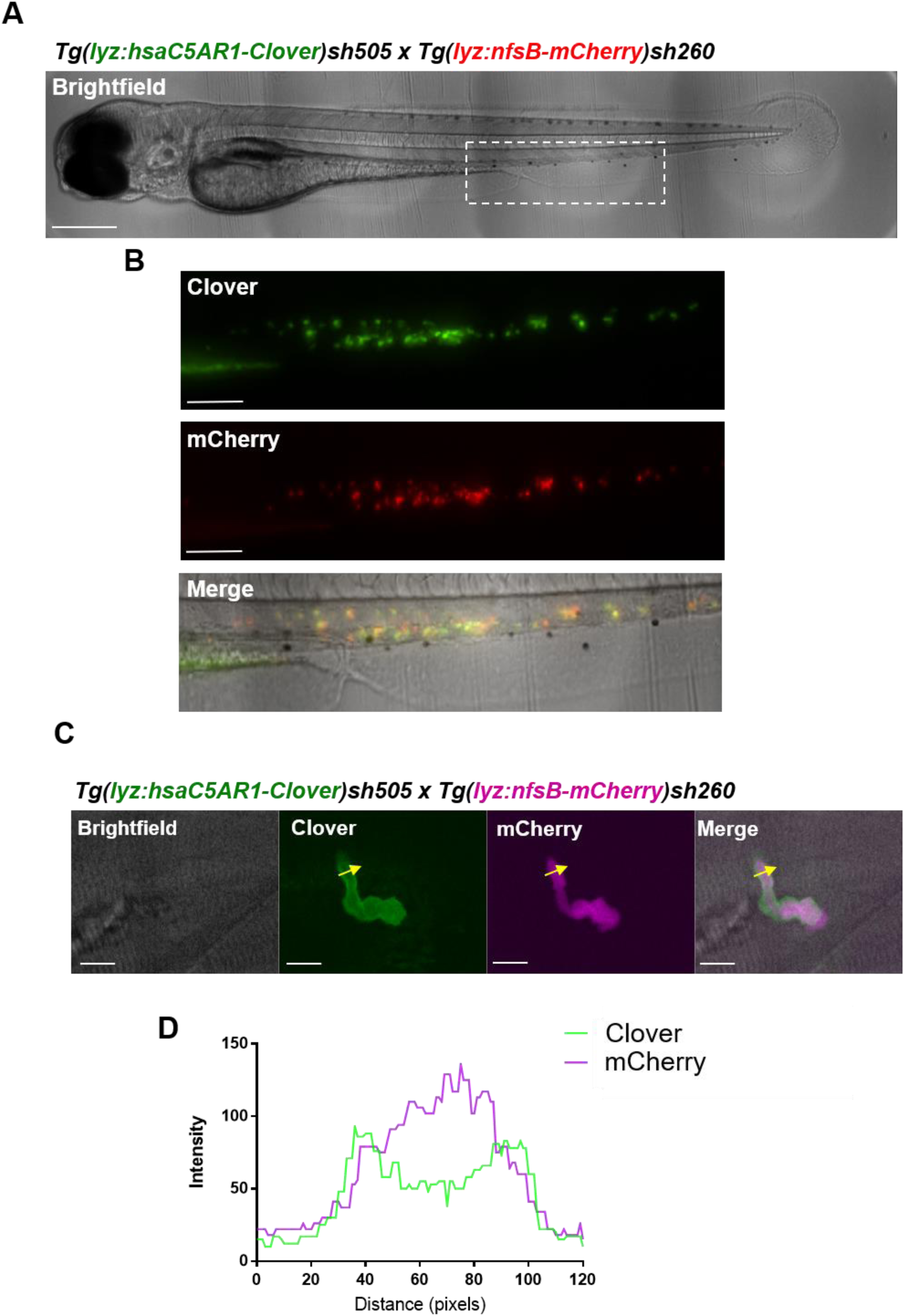
Generation of a transgenic zebrafish expressing human C5AR1-Clover. **A)** A double-transgenic *Tg(lyz:hsaC5AR1-Clover)sh505; Tg(lyz:nfsB-mCherry)sh260* larva at 3dpf. Dashed white box indicates the enlarged region shown in (B), scale bar = 250µm. **B)** Microscopy analysis of neutrophil-specific mCherry (red) and Clover (green) transgene expression in the enlarged view of the caudal haematopoietic tissue shown in (A), scale bar = 100µm. **C)** Close-up of Clover (green) and mCherry (purple) expression in a double-transgenic neutrophil in the caudal haematopoietic tissue of a 3dpf larva, scale bar = 10µm. **D)** Line intensity profile of the fluorescent signal of Clover (green) and mCherry (purple) across the yellow arrow shown in (C).

### Human C5AR1 is functional in humanised zebrafish

Next, we investigated functional activity of human C5AR1 in the transgenic zebrafish line by assessing neutrophil migration to recombinant dreC5a and hsaC5a injected into the otic vesicles, two sac-like invaginations in the head of the fish that are a preferred site for assessing phagocyte migration (*20*). In non-humanised *lyz:*nfsB-mCherry fish, dreC5a injection resulted in migration of neutrophils to the injection site, as expected due to endogenous receptor function, while hsaC5a injection did not induce neutrophil migration (**Fig. 3A,B**). In contrast, neutrophils expressing the human *C5AR1* transgene displayed robust migration towards the site of hsaC5a injection, showing that C5AR1 acts as a functional C5a receptor in zebrafish neutrophils that is able to direct neutrophil migration.

**Figure 3.**
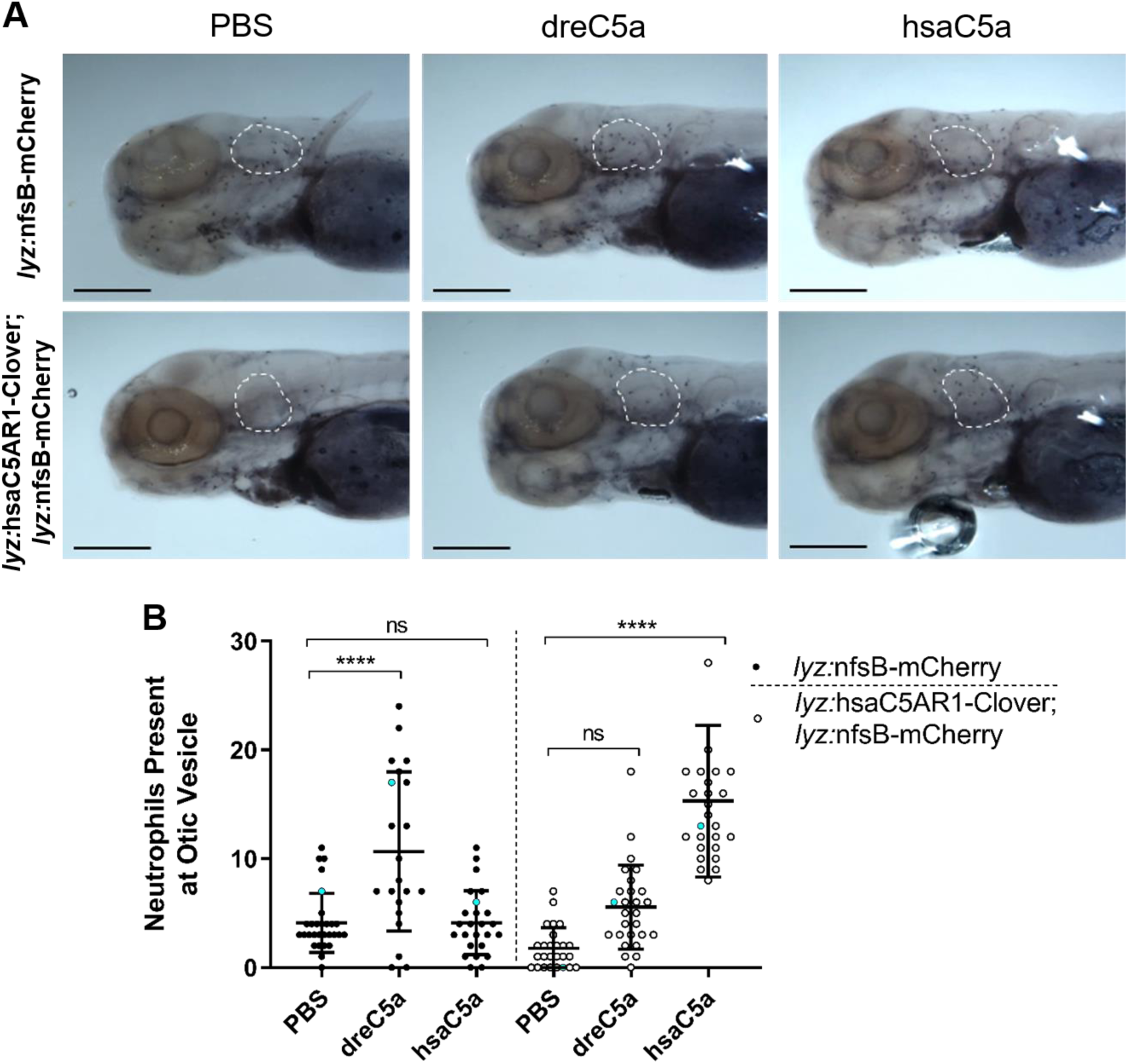
Human C5AR1 confers sensitivity to hsaC5a in humanised zebrafish. **A)** Neutrophil migration was assessed in non-humanised (*lyz:*nfsB-mCherry) and humanised (*lyz:*hsaC5AR1-Clover; *lyz:*nfsB-mCherry) zebrafish larvae at 3dpf following injection with a PBS vehicle control, recombinant zebrafish C5a (dreC5a) or human C5a (hsaC5a) into the otic vesicle. Four hours post injection (hpi), larvae were fixed in 4% paraformaldehyde and stained with Sudan Black B to detect neutrophils; scale bar = 200µm. **B)** Numbers of neutrophils present at the otic vesicle at 4hpi in zebrafish treated as in (A), blue points denote the representative images in (A). Error bars shown are mean ± SD (n=22-26 individual animals over two independent experiments); groups were analysed using a two-way ANOVA and adjusted using Bonferroni’s multiple comparisons test. ns, not significant; ****, *p*<0.0001.

### Humanised zebrafish neutrophils are targeted by PVL and HlgCB *in vivo*

Having established that human *C5AR1* is expressed as a functional receptor on the surface of zebrafish neutrophils, we next investigated whether *C5AR1*-expressing neutrophils are targeted by the C5AR1-targeting *S. aureus* toxins PVL and HlgCB *in vivo*. To this end, the community-acquired MRSA strain USA300 was injected with or without recombinant PVL into the otic vesicle of wild type or C5AR1-transgenic larvae, and the number of neutrophils present at the injection site was determined 4 hours later. Whereas injection of USA300 alone resulted in similar numbers of neutrophils at the injection site in both C5AR1 negative and positive larvae, the addition of PVL significantly reduced neutrophil numbers specifically in C5AR1-expressing larvae, while not affecting neutrophil migration in C5AR1-negative fish (**Fig. 4A,B**). Similarly, we observed a reduced number of neutrophils at the injection site in humanised larvae injected with USA300 together with HlgCB (**Fig. 4C,D**). These results showed that *C5AR1* expression sensitises zebrafish neutrophils to PVL and HlgCB *in vivo*.

**Figure 4.**
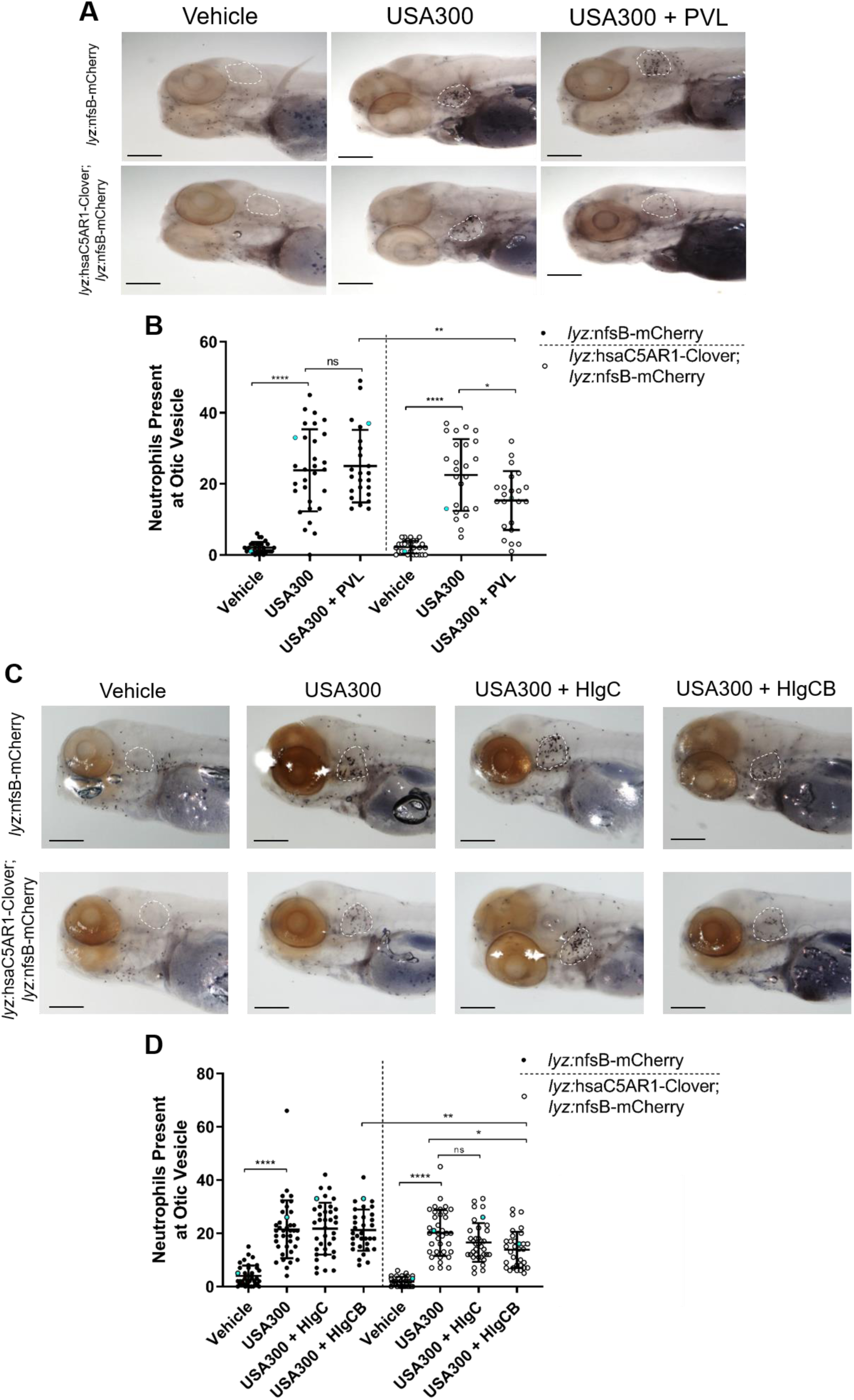
Neutrophils of C5AR1-expressing zebrafish are susceptible to *S. aureus* toxins PVL and HlgCB. **A)** Representative images of neutrophil abundance in zebrafish larvae at 3 dpf that were separated into non-humanised (*lyz:*nfsB-mCherry only) and humanised (*lyz:*hsaC5AR1-Clover; *lyz:*nfsB-mCherry) groups and injected into the otic vesicle with a vehicle control or ∼3,500cfu of *S. aureus* USA300 with or without 30.3 µg/ml PVL. The white outline indicates the otic vesicle; scale bar = 200µm. **B)** Number of neutrophils present at the otic vesicle at 4hpi, blue points denote the representative images in (A). **C)** Representative images of neutrophil abundance in zebrafish larvae injected into the otic vesicle as in (A) with a vehicle control, or ∼3,500 *S. aureus* USA300 with or without 16.7 µg/ml HlgCB or HlgC, as indicated. **D)** Number of neutrophils present at the otic vesicle at 4hpi, blue points denote the representative images in (C). Error bars shown are mean ± SD, groups were analysed using a two-way ANOVA and adjusted using Bonferroni’s multiple comparisons test. (B) n=22-26 over two independent experiments and (D) n=32-41 over three independent experiments. ns, not significant; *, *p*<0.05; **, *p*<0.01; ****, *p*<0.0001. Scale bars = 200µm.

The S-subunit of HlgCB (HlgC) has been reported to competitively inhibit C5AR1 signalling at low concentrations (*8, 9*). However, HlgC-only injection did not significantly affect neutrophil abundance at the injection site, suggesting that the reduced neutrophil counts observed in the presence of HlgCB injection are due to pore-forming activity of HlgCB and cell lysis rather than inhibition of C5aR signalling by HlgC alone (**Fig. 4C,D**). In conclusion, our data indicated that human *C5AR1* expression in zebrafish neutrophils conferred sensitivity to the *S. aureus* toxins PVL and HlgCB, and showed that the presence of these toxins reduces neutrophil numbers at the sites of infection *in vivo*.

### Zebrafish expressing human C5AR1 are more susceptible to *S. aureus* infection

Given that human C5AR1 acted as a functional receptor that was targeted by the *S. aureus* toxins PVL and HlgCB in our zebrafish model, we next sought to determine whether neutrophil-specific expression of C5AR1 increases the susceptibility of humanised fish to staphylococcal infection. To investigate this, we first separated the fish into non-humanised (*lyz:*nfsB-mCherry only) and humanised (*lyz:*hsaC5AR1-Clover; *lyz:*nfsB-mCherry) groups and injected the community-acquired MRSA strain USA300 into the circulation valley of the fish at 30hpf according to previously published methods (*21*). In this model, macrophages are able to clear *S. aureus* from the fish circulation, so to specifically study the effect of neutrophils, we silenced *irf8* expression using an *irf8* morpholino to alter zebrafish haematopoiesis and favour the differentiation of neutrophils over macrophages (*22*). In this way, we observed significantly higher mortality following staphylococcal infection for the C5AR1-positive zebrafish compared to the C5AR1-negative fish, both when infected at 30hpf or 50hpf (**Fig. 5A,B**). This demonstrates that expression of human *C5AR1* in zebrafish neutrophils enhances susceptibility to staphylococcal infection and suggests that the C5aR-targeting toxins PVL and HlgCB play crucial roles in determining the severity of *S. aureus* infection.

**Figure 5.**
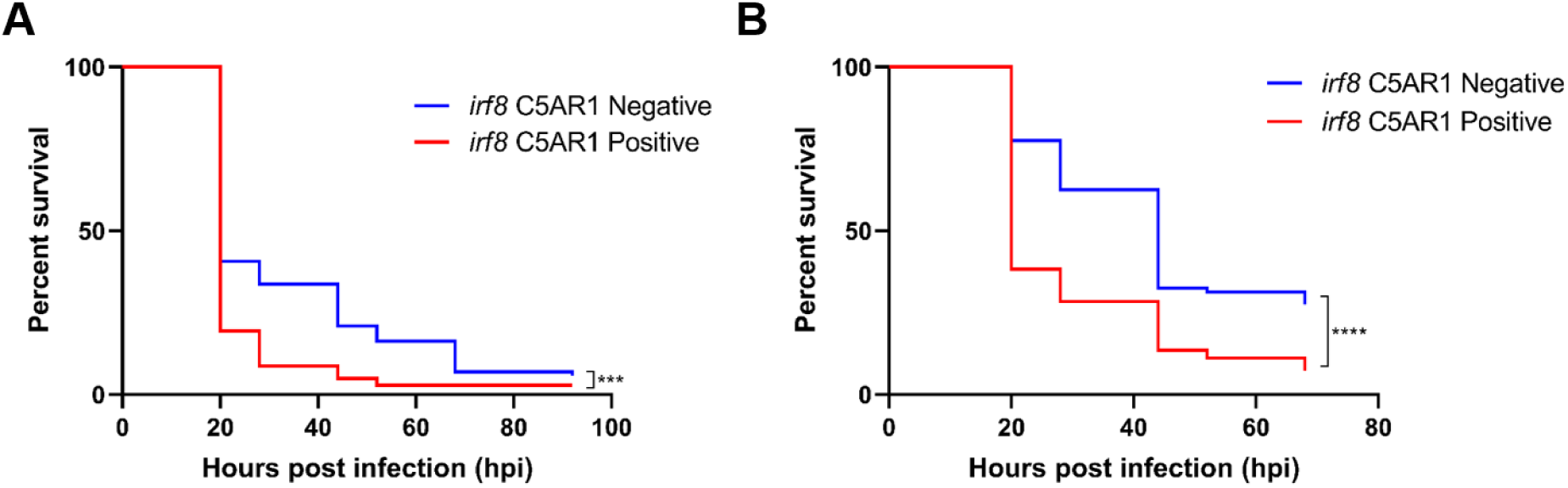
Humanised neutrophil-replete zebrafish are more susceptible to staphylococcal infection. Neutrophil-replete zebrafish were generated by injecting an *irf8* morpholino at the single-cell stage, silencing *irf8* expression. Zebrafish larvae were then separated into non-humanised (*lyz:*nfsB-mCherry only) and humanised (*lyz:*hsaC5AR1-Clover; *lyz:*nfsB-mCherry) groups and infected with **A)** 30hpf (∼700cfu), or **B)** 50hpf (∼2,000cfu). Survival was monitored up to 96 hrs post infection and data was analysed using a Log-rank Mantel-Cox test. (A) n= 80-81 over three independent experiments; ***, *p*<0.001. (B) n=86-103 over three independent experiments; ****, *p*<0.0001.

## Discussion

Complete understanding of staphylococcal infection requires a thorough characterisation of its virulence factors and an appreciation of their physiological relevance and synergistic role during natural infection. Due to a lack of appropriate animal models, human-specific virulence factors have been difficult to adequately study *in vivo*, creating a gap in our understanding that has disrupted the experimental validation of effective *S. aureus* vaccine candidates and novel therapeutic approaches. In this study, we addressed this problem by generating a humanised zebrafish model that allowed us to study the important contributions of human-specific *S. aureus* toxins PVL and HlgCB to infection-related mortality in a relevant *in vivo* system.

*S. aureus* expresses three virulence factors that target the human complement receptor C5aR, which is highly expressed on the surface of human neutrophils. Although components from all arms of the complement system (classical, alternative, lectin) have been found in larval zebrafish and are known to confer early humoral immunity to the embryo via maternal transfer (*23–25*), the zebrafish complement system including C5aR has barely been characterised at the functional level. To facilitate our studies of the C5aR-targeting virulence factors, we first established functionality of the C5a-C5aR axis in zebrafish. By expressing the zebrafish C5aR on U937 cells we showed that dreC5aR ligation by recombinant zebrafish C5a or zymosan-activated zebrafish serum resulted in similar Ca^2+^ mobilization as observed upon activation of the human receptor. Furthermore, we observed neutrophil migration toward the sites of dreC5a injection *in vivo*. Together, these results suggest that the C5a-C5aR axis is functional in zebrafish and, analogous to the human situation, is involved in directing neutrophil migration towards invasive pathogens. Interestingly, both human and zebrafish C5a displayed strict species specificity and were not interchangeable with one another. We further found that, in agreement with studies in other species (*8–10*), *S. aureus* virulence factors CHIPS, PVL, and HlgCB were ineffective against the zebrafish C5aR receptor, corroborating their strict human specificity.

The redundant targeting of human neutrophils by multiple *S. aureus* virulence factors suggests that inhibition of neutrophil function is an important contributor to *S. aureus* infection success. Neutrophils play an essential role in protecting the body from acute bacterial infection, and play a prominent role in clearing *S. aureus* infections (*7*). Importantly, the *S. aureus* toxins are thought to exacerbate infection-associated morbidity and mortality (*26*), however, due to the well-characterised human specificity of the interaction between C5aR and these toxins, their contribution to infection *in vivo* has remained elusive. The introduction of the human C5aR into zebrafish neutrophils allowed us to study the contribution of PVL and HlgCB-mediated targeting of the C5aR on neutrophils during natural infection and with high-throughput. In this way, we found that the actions of both toxins result in reduced neutrophil presence at the site of infection and increased infection-associated mortality. The presence of HlgC alone, which was recently shown to inhibit neutrophil chemotaxis (*9*), did not affect neutrophil numbers, suggesting that it is the cytotoxic activity of these toxins that leads to reduced neutrophil numbers in these fish, and not a blockade of neutrophil migration.

In humans, neutrophils play a critical role in containing and clearing invasive *S. aureus* infections. Uniquely, in zebrafish systemic infection is primarily controlled by macrophages, while neutrophils are recruited almost solely to surface-associated infections (*27*). To mimic human infections more closely and to study the role of neutrophils specifically, we utilised the *irf8* morpholino, which causes macrophages to undergo a cell-fate switch by suppression of the irf8 transcription factor (*22*). This depletes macrophages and promotes differentiation into neutrophils, producing neutrophil-replete zebrafish larvae that can be used to assess the role of toxin-mediated virulence in a way that is more representative of a human *S. aureus* infection. Neutrophil-replete zebrafish expressing the human C5aR were significantly more susceptible to *S. aureus* infection compared with C5AR1-negative siblings both in fish infected from 30 hours post-fertilisation (hpf) and also from 50 hours post-fertilisation. In both cases, *S. aureus* infection produces significant mortality from as early as 20 hours post-infection and is significantly enhanced by the expression of the human C5aR. Interestingly, fish infected at 50hpf appear to be twice as susceptible to those infected at 30hpf, accounting for a 40% and 20% increase in mortality respectively compared with C5AR1-negative siblings. This could be the result of an increased number of neutrophils at 50hpf compared with 30hpf (*14*), suggesting that mortality is further enhanced by the direct interaction between hC5aR-expressing neutrophils and human-adapted virulence factors.

Our work shows that selective humanisation of the zebrafish is a powerful approach towards identifying the contribution of host-restricted virulence to bacterial infection and pathogenesis. Interestingly, several other *S. aureus* virulence factors target GPCRs. For example, CXCR2 is targeted by SSL5, Staphopain A, and Leukotoxin ED (*28–30*) and the formyl peptide receptors (FPR1, FPR2) are targeted by CHIPS, FLIPr and SSL13 (*31–33*). We anticipate that more extensive humanisation of these and other receptors in zebrafish will lead to infection models that incorporate a multitude of human-specific virulence factors and even more closely resemble human infections, permitting detailed investigation of the interplay and relative importance of these virulence factors at the difference stages of infection. Our ultimate goal is to only marginally alter the endogenous zebrafish receptors at specific amino acids to minimally interfere with the zebrafish physiology. Our detailed knowledge of the receptor interaction sites will be harnessed to design minimally humanised receptors that gain susceptibility to the *S. aureus* virulence factors while maintaining *in vivo* functionality. Unfortunately, we were so far unable to generate a partially humanised zebrafish C5aR that displayed sensitivity to the pore-forming toxins and was still functionally expressed at the cell surface. Amino-acid substitutions in the extracellular loops of the receptor quickly abolished surface expression and thereby led to non-functional receptors, forcing us to introduce the entire human receptor into zebrafish neutrophils. It is promising however that only three amino-acid changes appeared sufficient to confer CHIPS-mediated inhibition to an otherwise functional receptor *in vitro*.

In conclusion, we show that humanised zebrafish are a powerful tool to study the contribution of human-specific *S. aureus* virulence factors to infection outcome *in vivo*. Importantly, we believe that this model provides a starting point that can be further developed to incorporate additional human-specific virus-host interactions, functioning as an improved, expandable, and translatable platform to accurately assess the efficacy of *S. aureus*-targeted therapeutic interventions.

## Materials & Methods

### Cells, Lentiviral transductions

Human monocytic U937 cells and HEK293T cells were obtained from ATCC (American Type Culture Collection) and grown in RPMI or DMEM medium, respectively, supplemented with glutamine, penicillin/streptomycin and 10% FBS. C5AR1 (CD88; NM_001736) and C5aR (XM_005159274) were cloned into a dual promoter lentiviral vector (BIC-PGK-Zeo-T2a-mAmetrine; RP172) described elsewhere (*34*). This vector expresses the cloned transgene from an EF1A promoter as well as the fluorescent protein mAmetrine and the selection marker ZeoR from the PGK promoter. Third generation lentiviral particles were produced in HEK293T cells following standard lentivirus production methods. Spin infection of U937 cells was performed by adding 100 µl virus supernatant with 8 µg/mL polybrene to 50.000 cells and spinning at 1000*g* for 2 h at 33 °C. Transduced cell lines were selected to high purity (>95%) by selection with 400 µg/ml Zeocin starting two days post transduction, and transgene expression was confirmed by flow cytometry using a mouse anti-FLAG M2 antibody (F1804, Sigma-Aldrich) together with an APC or PE-conjugated secondary anti-mouse antibody (Jackson) and acquisition on a FACSCantoII (BD Bioscience) cytometer. For expression of the zebrafish C5a receptor the full-length mRNA coding sequence was used (XM_005159274.1).

### C5aR-signalling assay

U937 cells were incubated with 2 mM Fluo-3AM (Thermo Fisher) in RPMI with 0.05% human serum albumin (HSA) at room temperature under constant agitation for 10 min, then washed and suspended to 3 million cells per mL in RPMI/0.05% HSA followed by data acquisition on a FACSVERSE (BD Biosciences). Basal fluorescence level for each sample was determined during the first ten seconds, followed by addition of the stimulus while continuing the acquisition to measure signalling-induced cytosolic Ca^2+^ release by increased Fluo-3AM fluorescence.

### Flow cytometry analysis of C5a binding

Binding of FITC-labelled hsaC5a (human C5a) or dreC5a (zebrafish C5a) was determined by incubating U937 cells with 10 μg/mL FITC-C5a in RPMI supplemented with 0.05% HSA for 30 minutes on ice. After washing the samples were analysed on a FACSVERSE (BD Biosciences).

### Collection of zebrafish serum

Zebrafish blood was kindly supplied by Dr. Astrid van der Sar (Amsterdam University Medical Centre) and serum collection was performed following a previously published protocol (*35*). Subsequently, 10% serum in HEPES buffer was incubated with zymosan for 30 minutes at 37 °C to activate the alternative complement pathway that results in C5a generation. The activated serum was centrifuged at 10,000 rpm and the supernatant containing the anaphylatoxins was stored at −80°C.

### Cell permeability assays

Cells were resuspended in 100 μL RPMI/0.05% HSA and incubated with for 30 min at 37 °C with 10 μg/mL PVL or HlgCB (as PVL and HlgCB are two-component toxins, equimolar concentrations of polyhistidine-tagged LukS-PV, LukF-PV, HlgC and HlgB were used). Cells were then stained with 1 μg/mL 4′,6-diamidino-2-phenylindole (DAPI; Molecular Probes/Thermo Fisher) and analysed on a FACSVERSE (BD Biosciences). Pore formation was defined as the percentage of cells positive to DAPI staining.

### Recombinant protein production and FITC labelling

LukS-PV, LukF-PV, HlgC and HlgB were cloned and expressed as previously described (*8, 9, 36*). From the coding sequence of zebrafish C5 (XM_001919191.5) we identified a predicted C5a cleavage product (KFE DKA QKY GAF REY CLS GTR SSP TLE TCK DRA NRV TLP NKK TRR DYE KEK YCR LAF EQC CVF AKD LRK E) and included nine additional amino acids from an alignment with human C5 (NAE NII LSR). This sequence was codon-optimised for expression in *E. coli* K-12 and ordered as a gBlock (Integrated DNA Technologies). This was then ligated into the BamHI and NotI sites of the modified expression vector pRSETB (Invitrogen Life Technologies), containing a cleavable N-terminal poly-histidine tag and three glycines (6xHis-TEV-GGG-dreC5a) or a non-cleavable N-terminal polyhistidine tag. Zebrafish C5a was expressed in *E. coli* strain Rosetta-gami(DE3)pLysS (Novagen; Merck Biosciences). Following cell lysis with 10 µg/mL lysozyme and three freeze– thaw sonication cycles in 20 mM sodium phosphate (pH 7.8), the His-tagged proteins were purified using nickel-affinity chromatography (HiTrap chelating, HP; GE Healthcare) with an imidazole gradient (10-250 mM; Sigma-Aldrich). Purified protein was stored in PBS at −20 °C. Subsequently, the polyhistidine tag of 6xHis-TEV-dreC5a was removed by incubation with TEV protease (Thermo Fisher Scientific) and the protein was FITC-labelled at the N-terminus using the sortagging method (*37*). CHIPS protein was purified using previously published methods (*32, 38*).

### Zebrafish Husbandry

Zebrafish (*Danio rerio*) were raised and maintained according to standard protocols (*39*) in UK Home Office-approved aquaria at the Bateson Centre, University of Sheffield, and kept under a 14/10 hour light/dark regime at 28°C.

### Creation of *Tg(lyz:hsaC5AR1-Clover)sh505* zebrafish

The plasmid used for introducing the transgene into the zebrafish genome (pDestTol2CG2 *lyz:*hsaC5AR1-Clover *cmlc2*:eGFP) was created by Gateway cloning (*19*). The *C5AR1* gene was PCR amplified from the pIRES-C5AR1 plasmid with a truncated stop codon to allow C’ terminal fusion of clover and ligated into the middle-entry clone vector pDONR221. The final construct was created by an LR reaction combining a 5’ vector containing the *lyz* promoter, the middle entry vector pDONR221 C5AR1-Clover, a 3’ vector containing the Clover fluorophore, and the destination vector pDestTol2CG2. To induce transgenesis, plasmid DNA of pDestTol2CG2 *lyz:*hsaC5AR1-Clover *cmlc2*:eGFP was injected into zebrafish embryos at the one-cell stage with 10ng/µl of Tol2 transposase mRNA, according to published protocols (*39*). At three days post-fertilisation, positive transgenic larvae were selected and raised to maturity, then screened for successful germline integration of the construct.

### Injection into the zebrafish otic vesicle

1% agarose dishes supplemented with E3 were cast using triangular moulds. Before injection, larvae were anaesthetised by immersion in 0.02% (w/v) Tricaine prior to transfer to the mounting dish. Larvae were then arranged laterally in rows. Excess media was then removed with a pipette to minimise movement during injection. Larvae were then injected from the dorsal side into the otic vesicle. Four hours after injection, larvae were fixed for 1 hour in 4% paraformaldehyde, and later stained with Sudan Black B to indicate neutrophils (performed according to published methods (*18*)). Proteins were injected at the highest available concentration, which were 10µM and 90µM for hsaC5a (human) and dreC5a (zebrafish) respectively, and 30.3µM and 16.7µM for PVL and HlgCB respectively.

### Systemic infection of zebrafish embryos and *irf8* knockdown

Zebrafish larvae at 30 or 50 hpf were microinjected into the circulation with bacteria as previously described (*21*). Briefly, anaesthetised larvae were embedded in 3% w/v methylcellulose and injected individually with 1 nl using microcapillary pipettes filled with the bacterial suspension of known concentration. Following infection, larvae were observed frequently up to 122 hpf and numbers of dead larvae recorded at each time point. Morpholino-modified antisense oligomer against and *irf8* (splice MO) (*22*) was injected using a method described previously (*21*).

### Bacterial Culture Preparation

To prepare a liquid overnight culture of *S. aureus*, 5ml of BHI broth medium (Oxoid) was inoculated with a colony of *S. aureus* strain USA300 and incubated at 37°C overnight with shaking. To prepare *S. aureus* for injection, 50ml of BHI media was inoculated with 500µl of overnight culture and incubated for roughly 2 hours at 37°C with shaking. The OD_600_ of each culture was measured and 40ml of the remaining culture harvested by centrifugation at 4500g for 15 minutes at 4°C. The pellet was then resuspended in a volume of PBS appropriate to the bacterial dose required. Once the pellets were resuspended they were then kept on ice until required.

### Statistics

All data were analysed in Prism 7.0 (GraphPad Software, San Diego, CA, USA). Comparisons between groups were performed using a two-way ANOVA with different multiple comparisons tests depending on whether the group was compared with a control (Dunnett’s test) or not (Bonferroni’s test). Significance was assumed at p<0.05.

## Supporting information

Supplementary Figure 1. Expression of receptor variants in U937 cells.

Supplementary Figure 2. Three amino-acid changes in the CHIPS binding site of dreC5aR confer susceptibility to inhibition by CHIPS.

Supplementary Figure 3. Expression of the human C5a receptor does not interfere with zebrafish haematopoiesis.

## Acknowledgements

### General

Thank you to C. Loynes and D. Drew for their training, advice and technical help. Also, we’d like to thank the Bateson Centre aquarium staff for all their help, A. M. van der Sar for facilitating the zebrafish blood collection, and the Wolfson Light Microscopy Facility.

### Funding

This work was supported by AMR cross-council funding from the MRC to the SHIELD consortium “Optimising Innate Host Defence to Combat Antimicrobial Resistance” (MRNO2995X/1), a Medical Research Council (MRC), United Kingdom, Programme Grant (MR/M004864/1) to S.A.R., and the University of Sheffield 2022 Futures programme via the Florey Institute. T.K.P. was supported by an individual Marie Curie fellowship (PIEF-GA-2013-625975).

### Author Contributions

K.D.B and M.v.G performed experiments with assistance from T.K.P, N.V.O, N.W.M.d.J, and J.K.; S.A.R, J.A.G.v.S and S.J.F conceived the study and designed experiments. T.K.P performed survival experiments on *irf8* morphants. N.V.O generated the *Tg(lyz:nfsB-mCherry)sh260* line. K.D.B, M.v.G and S.A.R wrote the manuscript with significant input from all authors.

### Competing Interests

The authors declare no conflicts of interest.

### Data and Materials Availability

The raw data for this manuscript is available at Mendeley Data, DOI: 10.17632/x55r5h6j4g.2.

## Abbreviations

BHI: Brain-Heart Infusion
C5aR: C5a receptor
C5AR1: Human C5a receptor
c5ar1: Zebrafish C5a receptor
cfu: colony forming units
CHIPS: Chemotaxis inhibitory protein of *Staphylococcus*
dpf: days post fertilisation
dpi: days post injection
dreC5a: Zebrafish C5a
EV: Empty Vector
FLIPr: FPRL1 Inhibitory Protein
FPR: Formyl Peptide Receptor
GPCR: G-Protein Coupled Receptor
HlgCB: γ-Haemolysin CB
hpf: hours post fertilisation
hpi: hours post injection
hsaC5a: Human C5a
irf8: Interferon Regulatory Factor 8
MO: Morpholino
MRSA: Methicillin-Resistant *Staphylococcus aureus*
nl: Nanolitre
OD_600_: Optical Density at 600 nanometres
PBS: Phosphate-Buffered Saline
PVL: Panton-Valentine Leukocidin
SD: Standard Deviation
SSL: Staphylococcal Superantigen-Like.

## Supplementary Figures

**Supplementary Figure 1.**
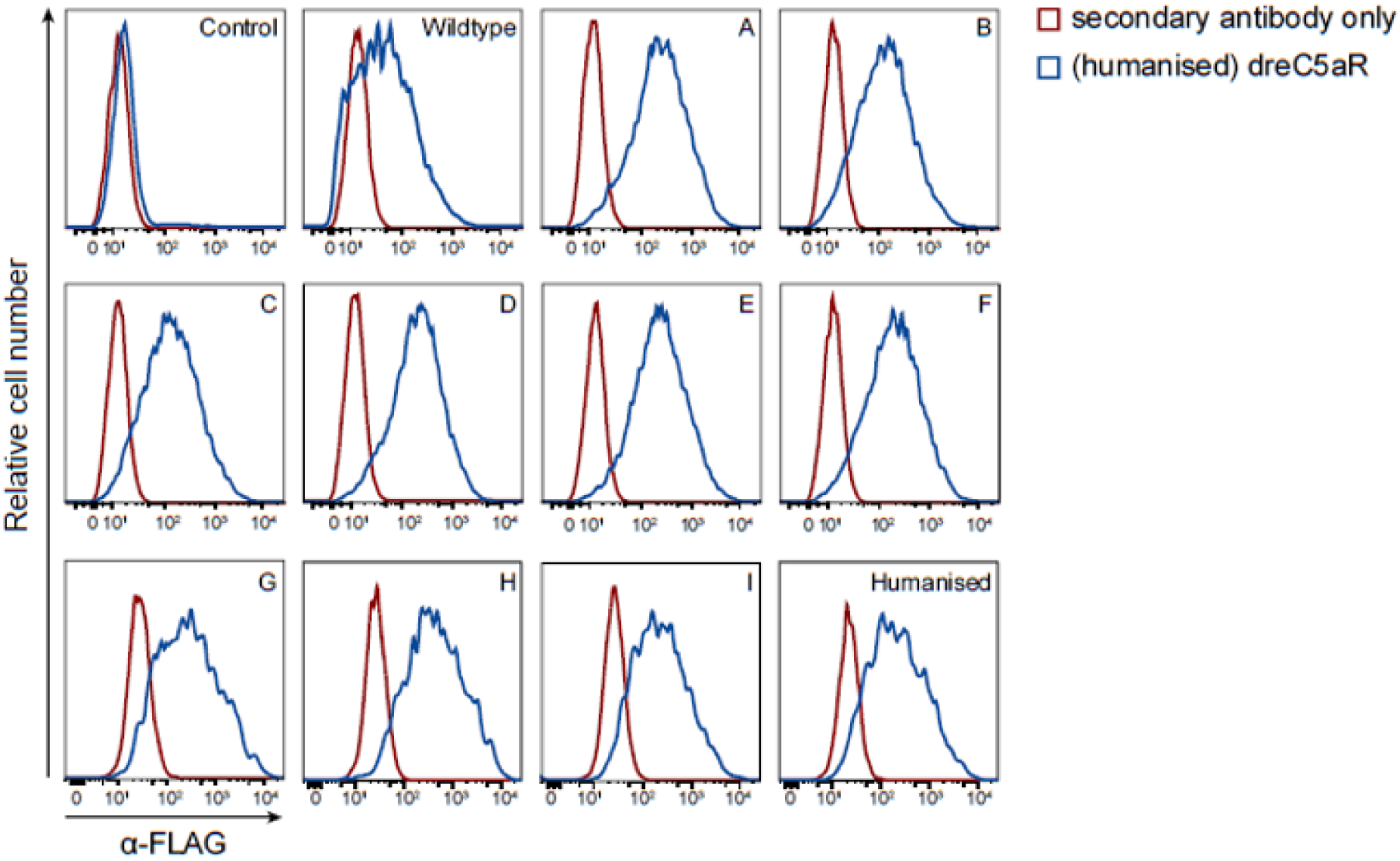
Expression of receptor variants in U937 cells. Expression of FLAG-tagged wild type, partially humanised dreC5aR receptor variants A-I, and dreC5aR with a fully humanised CHIPS binding site (‘humanised’) on transduced U937 cells was evaluated by flow cytometry following staining with an anti-FLAG antibody and an APC-conjugated secondary antibody (blue). Control, empty-vector transduced U937 cells. Staining without the primary antibody (red) was included as negative control.

**Supplementary Figure 2.**
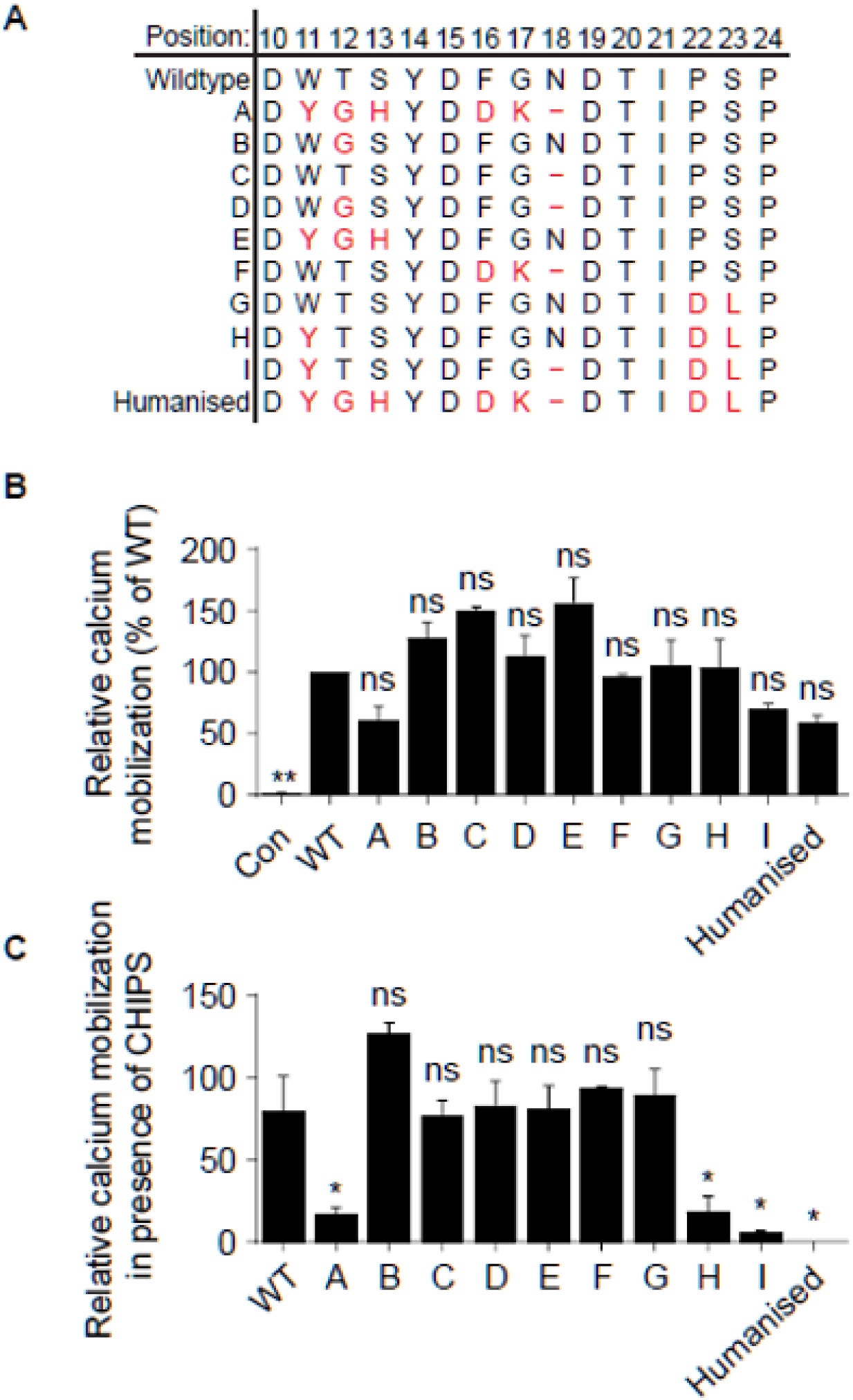
Three amino-acid changes in the CHIPS binding site of dreC5aR confer susceptibility to inhibition by CHIPS. **A)** Overview of the ten dreC5aR (A-I and fully humanised) variants with partially humanised N-termini tested; altered amino acids indicated in red, numbers refer to amino acid positions in the dreC5aR. **B)** Relative C5aR-signalling activity in U937 cells stably expressing an empty vector control (Con), the wild type dreC5aR (WT), or the dreC5aR variants described in (A); assessed by measuring calcium mobilisation by flow cytometry using the fluorescent probe Fluo-3AM following incubation with 3.3 μM recombinant His-dreC5a. Data is presented as relative fluorescent signal compared to the wild type U937-dreC5aR (WT, set at 100%). **C)** C5aR activity determined as in (B) following incubation with 3.3 μM recombinant His-dreC5a in the absence or presence of 10 μg/ml recombinant CHIPS; presented as percentage fluorescent signal of the CHIPS-treated sample relative to the CHIPS-negative control. All data are presented as means ± SD and were analysed using a two-way ANOVA with Dunnett’s multiple comparisons correction; *, *p*<0.05; **, *p*<0.01; ns, not significant.

**Supplementary Figure 3.**
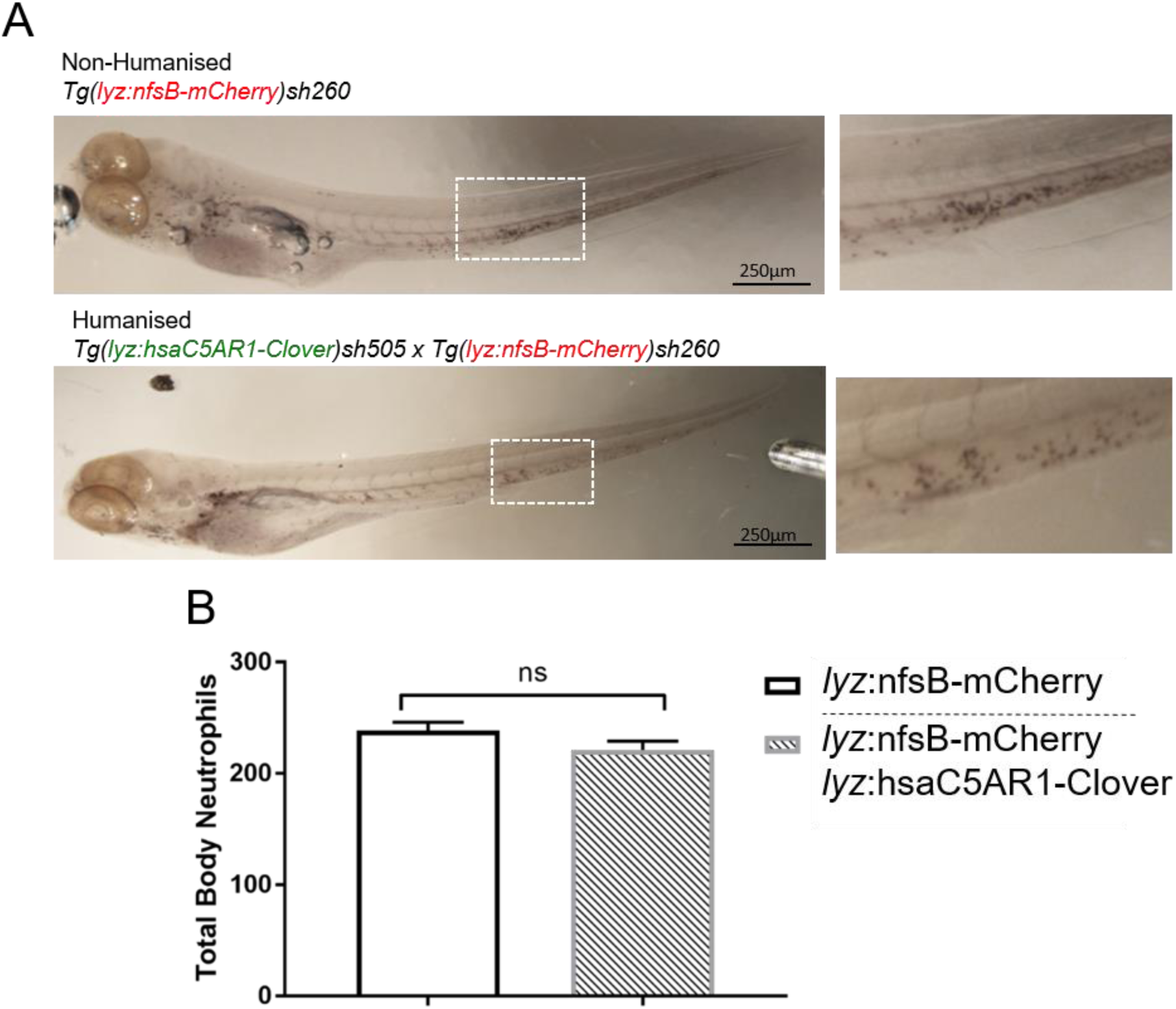
Expression of the human C5a receptor does not interfere with zebrafish haematopoiesis. **A)** 4dpf larvae from a *Tg(lyz:hsaC5AR1-Clover)sh505 x Tg(lyz:nfsB-mCherry)sh260* cross, separated into non-humanised (*lyz:*nfsB-mCherry only) and humanised (*lyz:*hsaC5AR1-Clover; *lyz:*nfsB-mCherry) groups and stained with Sudan Black B to detect neutrophils. The white dashed box indicates the enlarged view alongside. **B)** Total body neutrophil counts from both groups. Values shown are mean ± SEM (n=50 over two experiments); groups were analysed using an unpaired *t*-test (two-tailed). ns, not significant.

