## Supplementary figures and images for "Human-specific staphylococcal virulence factors enhance pathogenicity in a humanised zebrafish C5a receptor model"

### Supplementary Figure 1. Expression of receptor variants in U937 cells.

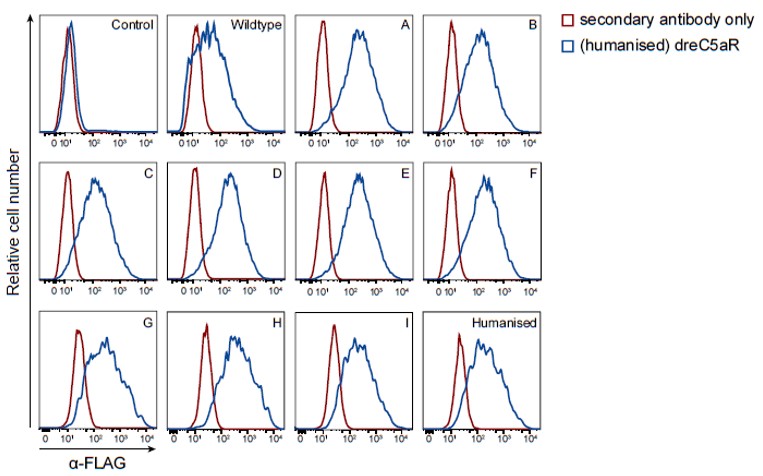

### Supplementary Figure 2. Three amino-acid changes in the CHIPS binding site of dreC5aR confer susceptibility to inhibition by CHIPS.

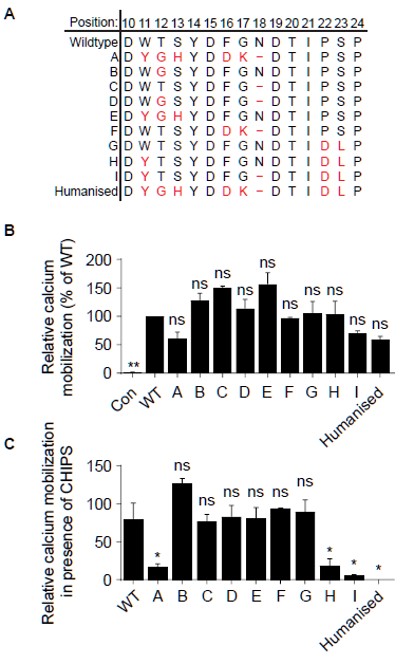

### Supplementary Figure 3. Expression of the human C5a receptor does not interfere with zebrafish haematopoiesis.

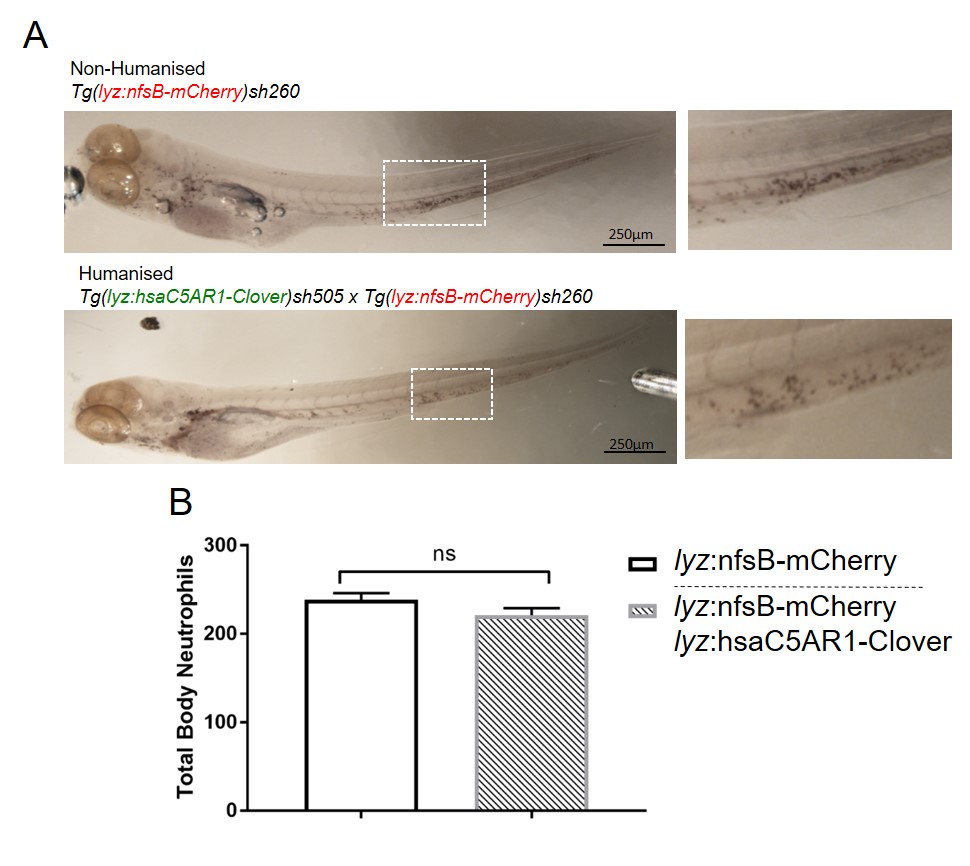
